# Non-canonical circadian oscillations in *Drosophila* S2 cells drive gene-expression cycles coupled to metabolic oscillations

**DOI:** 10.1101/191338

**Authors:** Guillaume Rey, Nikolay B. Milev, Utham K. Valekunja, Ratnasekhar Ch, Sandipan Ray, Mariana Silva Dos Santos, Andras D. Nagy, Robin Antrobus, James I. MacRae, Akhilesh B. Reddy

**Affiliations:** The Francis Crick Institute, 1 Midland Road, London NW1 1AT, UK; UCL Institute of Neurology, Queen Square, London WC1N 3BG, UK; Cambridge Institute for Medical Research (CIMR), Wellcome Trust/MRC Building, Addenbrooke’s Hospital, Cambridge CB2 0XY, UK

## Abstract

Circadian rhythms are cell-autonomous biological oscillations with a period of about 24 hours. Current models propose that transcriptional feedback loops are the principal mechanism for the generation of circadian oscillations. In these models, *Drosophila* S2 cells are generally regarded as ‘non-rhythmic’ cells, as they do not express several canonical circadian components. Using an unbiased multi-omics approach, we made the surprising discovery that *Drosophila* S2 cells do in fact display widespread daily rhythms. Transcriptomics and proteomics analyses revealed that hundreds of genes and their products are rhythmically expressed in a 24-hour cycle. Metabolomics analyses extended these findings and illustrated that central carbon metabolism and amino acid metabolism are the main pathways regulated in a rhythmic fashion. We thus demonstrate that daily genome-wide oscillations, coupled to metabolic cycles, take place in eukaryotic cells without the contribution of known circadian regulators.

## Introduction

Circadian rhythms are ∼24-hour oscillations found in virtually all aerobic life forms (1). In multi-cellular organisms, such timing mechanisms enable the temporal organisation of body physiology and behaviour, in synchrony with the alternation of day and night (2). In the fruit fly *Drosophila Melanogaster*, the recognized models of the circadian clock centre on the transcription factors CYCLE (CYC) and CLOCK (CLK), the homologs of BMAL1 and CLOCK in mammals. These control the transcription of several clock genes including *period* (*per*) and *timeless* (*tim*) (3). Only a subset of cells in *Drosophila* express these ‘clock’ components and are regarded as the principal pacemakers that drive daily activity rhythms. All other cells are not thought to have the capacity to generate 24-hour rhythms autonomously. This extends to *Drosophila* S2 cells, which are regarded as ‘non-rhythmic’ because they do not express several canonical circadian components, including PER, TIM or CLK (4, 5).

Several lines of evidence suggest that current models are not able to fully explain the extraordinary plasticity of circadian oscillations (6). Residual oscillations persist in several animal models with clock genes mutated (7-9) or constitutively expressed (10), suggesting that the canonical mechanisms are not sufficient to explain the emergence of circadian oscillations. In addition, circadian oscillations precede the expression of canonical circadian genes during embryonic differentiation (11, 12), suggesting that oscillatory behaviour is not fully dependent on clock genes. Moreover, the resilience of circadian oscillations to the reduction of RNA transcription (13) and the persistence of metabolic oscillations in complete absence of gene expression (14) further highlight that current models do not encompass the full range of oscillatory modes on the circadian time scale.

Therefore, we set out to investigate if a “clock-less” system such as *Drosophila S2* cells could exhibit circadian oscillations and would provide a novel paradigm to study fundamental mechanisms of circadian oscillations. To this end, we combined multiple “omics” technologies, including transcriptomics, proteomics and metabolomics, which identified several hundred transcripts and proteins rhythmically expressed in these cells. Importantly, metabolic processes were the main cellular function controlled by these non-canonical circadian oscillations in both transcriptomics and proteomics datasets. Metabolomics analyses extended these findings by showing that central carbon metabolism and amino acid metabolism were regulated in a rhythmic fashion, while integration of proteomics and metabolomics data revealed a strong coupling between protein and metabolite accumulation.

## Results

### Defining the daily transcriptome in *Drosophila* S2 cells

We first explored gene expression patterns in S2 cells using RNA Sequencing (RNA-Seq). To ensure that the cells were synchronized (’entrained ‘), we employed 24-hour temperature cycles. These have been shown to be efficient synchronisation signals for circadian oscillations (15). Following synchronization for a week, we sampled cells at 3-hour intervals in constant conditions (i.e. in the absence of any external temporal cues) (Figure 1A), and measured their circadian transcriptome by RNA-Seq (Figure S1A). Using a mixture model to define the sets of low and highly expressed transcripts (Figure S1B), we established that several clock genes including *clk*, *per* and *tim* were not expressed in S2 cells (Figure 1B), consistent with previous studies (4, 5). Moreover, four canonical clock genes that are expressed in S2 cells – *clockwork orange* (*cwo*), *cyc, par domain protein 1* (*pdp1)* and *vrille* (*vri*) – did not exhibit circadian rhythmicity (Figure S1C). This verified that any known circadian components were either absent or arrhythmic in this cell line.

**Figure 1.**
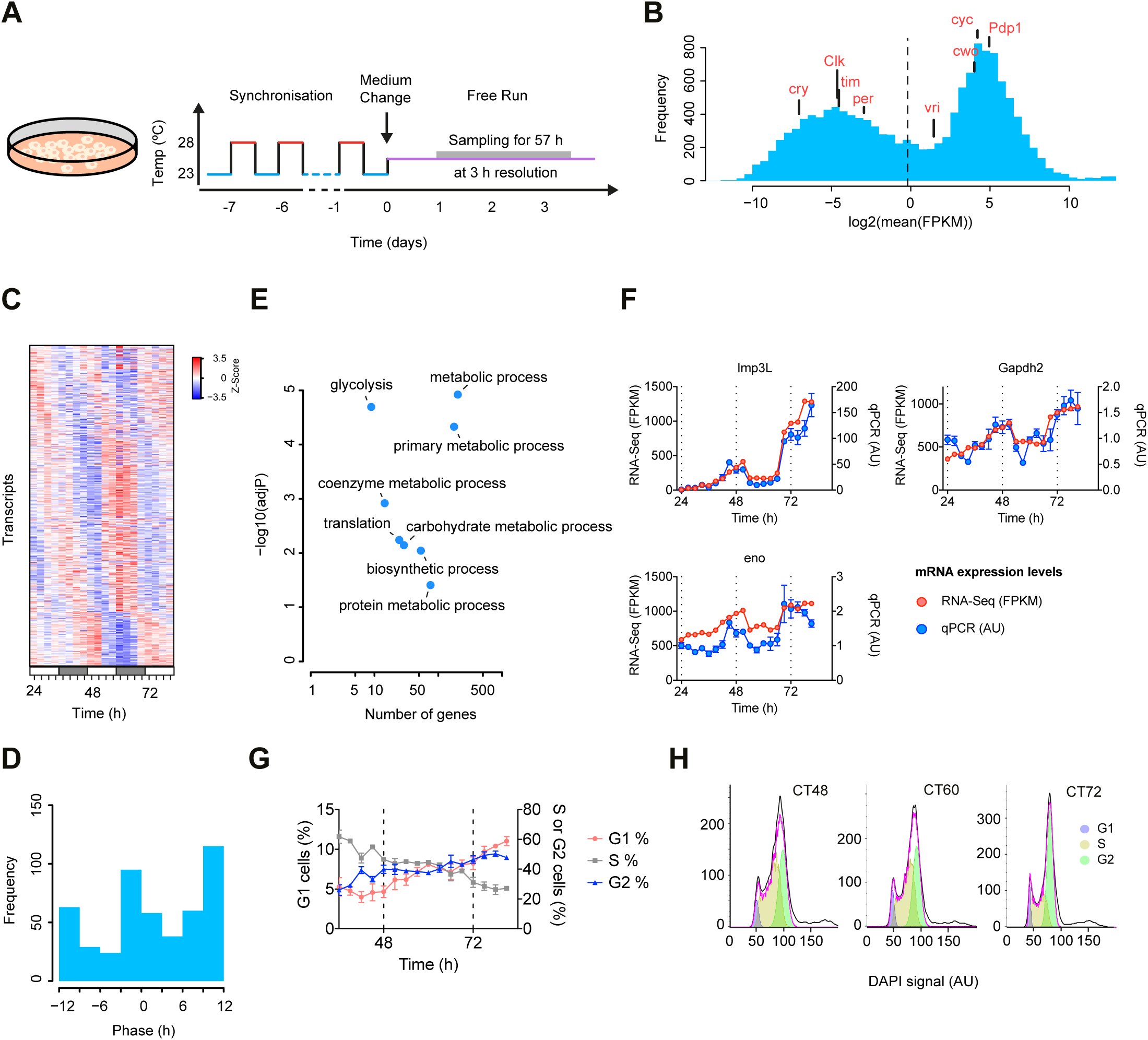
Transcriptional oscillations in *Drosophila* S2 cells. (A) Schematic showing the entrainment protocol used to synchronize *Drosophila* S2 cells. (B) Histogram showing the distribution of mean expression levels. The dashed vertical line represents the cut-off chosen to define the set of expressed transcripts. FPKM, Fragments Per Kilobase of transcript per Million mapped reads. (C) Heatmap showing the expression profiles of the 511 rhythmic transcripts (JTK-Cycle, p < 0.05 (D) Distribution of phase of the circadian transcripts shown in (C). (E) Scatterplot representation of Gene Ontology (GO) analysis of the rhythmic transcripts. (F) Validation of selected transcripts by quantitative PCR (qPCR) (n = 3, mean ± SEM). (G) Cell cycle analysis using flow cytometry showing the fraction of cells in G1, S and G2 phases along the time course experiment (n = 3, mean ± SEM). (H) Representative histograms of the DAPI signal at selected time points, with the curve fitting of cell cycle phases.

In contrast, the JTK-Cycle algorithm (16) detected more than 500 rhythmic transcripts with a period of approximately 24 h (Figure 1C-D; JTK-Cycle, adjusted p-value < 0.05), with peak phases at CT0 and CT12 (Figure 1D). Furthermore, we found a similar number of rhythmic transcripts using two alternative detection methods (RAIN (17) and ARSER (18)), with a highly significant overlap between the groups (Figure S1D; Fisher test on contingency table, p < 10^−16^ for every pairwise comparison). Thus, without any known ‘clock genes’ being rhythmic, 24-hour gene expression cycles were readily apparent in these cells.

Gene Ontology (GO) analyses revealed that rhythmic genes were enriched for protein biosynthesis and metabolic processes and, in particular, the glycolytic pathway (Figure 1E). To validate this finding, we measured the mRNA accumulation of three glycolytic genes – *lactate dehydrogenase* (*ImpL3*), *enolase* (*eno*) and *Glyceraldehyde 3-phosphate dehydrogenase 2* (*Gapdh2*) – using quantitative PCR (qPCR) and found strong agreement with our RNA-Seq data (Figure 1F).

Since S2 cells were actively dividing in culture, we investigated whether the cell cycle could be contributing to the rhythmic transcription that we observed. We therefore performed flow cytometry using DAPI staining to determine the cell cycle status of temperature-synchronized cells over two days. We did not see circadian variation in the proportion of cells in G1, S and G2 phases during our time course (Figure 1G-H), indicating that the phasing of the circadian and cell cycles is not significantly correlated. These results thus suggest that an uncharacterised mechanism, independent of canonical circadian genes or the cell cycle, is involved in the generation of 24-hour transcriptional oscillations in S2 cells.

### The daily proteome is enriched for abundant proteins with low amplitude oscillations

In addition to metabolic processes, we observed an overrepresentation of transcripts from protein translation and protein metabolic processes (Figure 1E). We therefore hypothesized that large-scale changes in protein levels may be occurring on a 24-hour timescale. We first quantified the total protein content over a time course and found that it displayed a circadian pattern (Figure S2A). Given this, we proceeded to determine whether specific proteins were rhythmically regulated using a proteomics protocol based on isobaric labelling of peptides using tandem mass tags (TMTs) (Figure 2A). Using this approach, we were able to reliably quantify 4759 proteins over 18 time points and found that 342 proteins were rhythmically expressed in this system (Figure 2B; JTK-Cycle, adjusted p-value < 0.05). We again cross-validated our rhythmicity analysis using two other detection methods and found a similar number of rhythmic proteins (Figure S2B; Fisher test, p < 10^−16^ for every pairwise comparison). As was the case for RNA transcripts, the distribution of phases displayed a biphasic pattern (Figure 2C). However, most proteins peaked at CT0, while the peak phase for transcripts was CT12 (Figure 1C-D). Together these results indicate that S2 cells generate 24-hour gene expression cycles in tandem with rhythmic regulation of protein translation of specific proteins.

**Figure 2.**
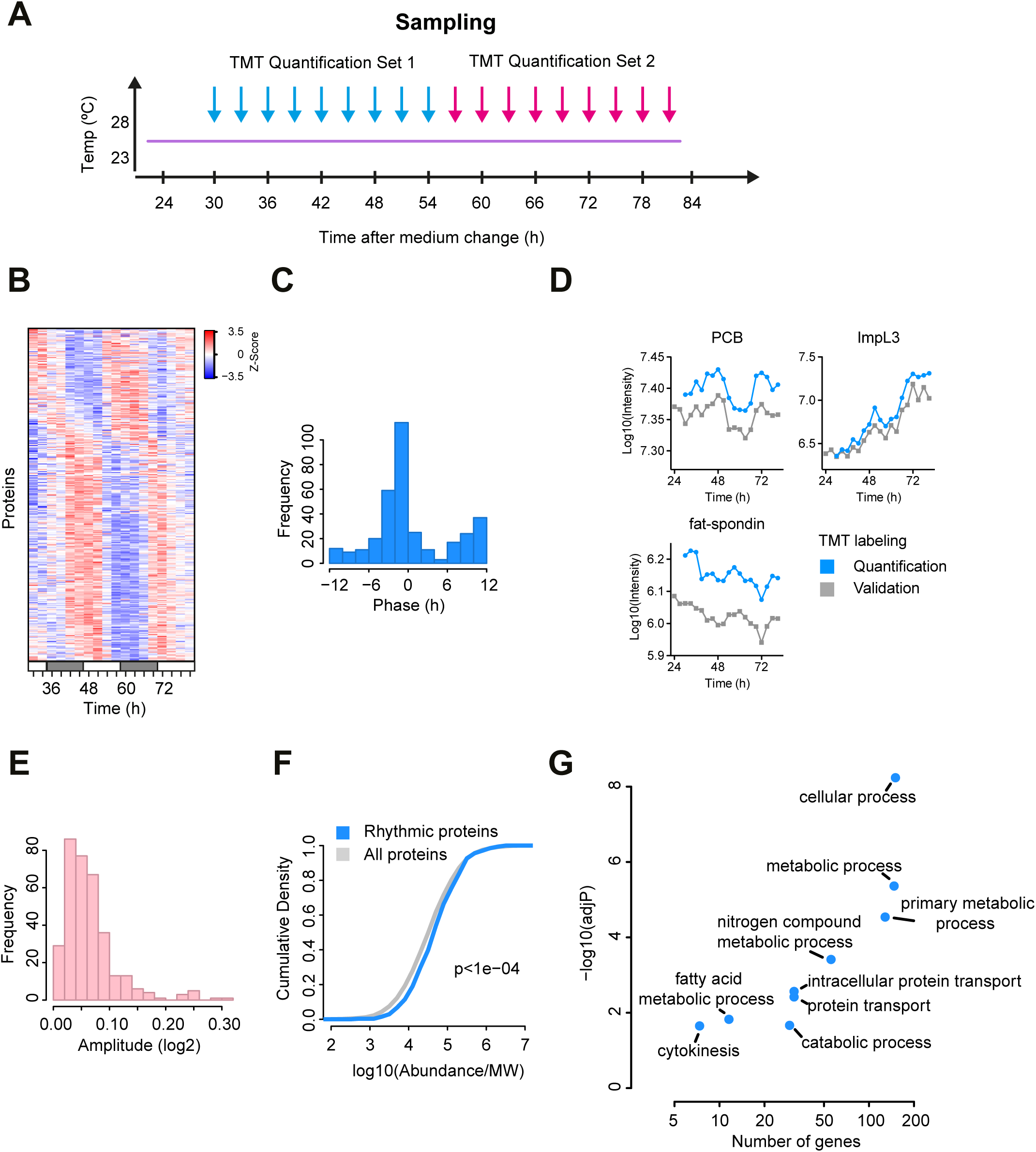
Proteome oscillations in *Drosophila* S2 cells. (A) Scheme showing the sample collection procedure and the labelling scheme. (B) Heatmap representation of the 342 rhythmic proteins (JTK-Cycle, p < 0.05). (C) Distribution of phase of the circadian proteins shown in (B). (D) Validation of TMT quantitation using an alternative method to label samples. (E) Distribution of amplitudes of the 342 circadian proteins. (F) Cumulative density of protein abundances of circadian proteins vs. all proteins (n = 4759 proteins quantified; Wilcoxon sum rank test, p < 10^−4^). (G) Scatterplot representation of the GO analysis of the rhythmic proteins.

We next assessed the reliability of our protein quantification using an alternative labelling strategy (Figure S2C). Correlation analysis between the temporal profiles generated by the two labelling strategies showed that our TMT-based protocol yields reproducible temporal profiles (Figure S2D). Examples of this included ImpL3, Pyruvate carboxylase (PCB) and fat-spondin, which displayed almost identical profiles using both quantification methods (Figure 2D). Moreover, the set of 342 circadian proteins showed similar profiles in the validation set (Figure S2E) and the overlap between circadian proteins in the quantification and validation sets was highly significant (Figure S2F) (Fisher test, p < 5x10^−4^). Similarly to previous studies in mammals (19, 20), the amplitude of protein oscillations was relatively low (Figure 2E). In addition, we observed that rhythmic proteins were enriched among the abundant proteins (Figure 2F). Since the latter tend to be more stable than less abundant ones (21), the low amplitude oscillations of rhythmic proteins may be caused by the enrichment of highly expressed proteins. Interestingly, GO analysis revealed that metabolic processes were particularly enriched among rhythmic proteins (Figure 2G), in a similar manner to our transcriptomics analyses (Figure 1E). Together, these results show that the circadian proteome in S2 cells is enriched for abundant proteins, likely due to the overrepresentation of metabolic enzymes, and identify cellular metabolism as a key process regulated in a circadian pattern.

### Different contributions of transcriptional and post-transcriptional mechanisms to the circadian proteome

Having determined the circadian transcriptome and proteome in S2 cells, we set out to understand how circadian transcription may contribute to rhythmic protein accumulation. First, we observed that the abundance of transcripts and proteins were highly correlated (Figure 3A), whereas there was little overlap of rhythmic transcripts and proteins (Figure 3B), suggesting that post-transcriptional mechanisms play an important role in shaping the circadian proteome. These results are in agreement with previous studies in mammalian systems (19, 20, 22). Nevertheless, we observed significant temporal correlations among RNA-protein pairs. Interestingly, the correlation between transcript and protein was indeed significantly higher for RNA-protein pairs when either the transcript or the protein (or both) was rhythmic (Figure 3C). This indicates that, even if RNA-protein pairs are rarely detected as both rhythmic, they nevertheless exhibit correlated temporal profiles.

**Figure 3.**
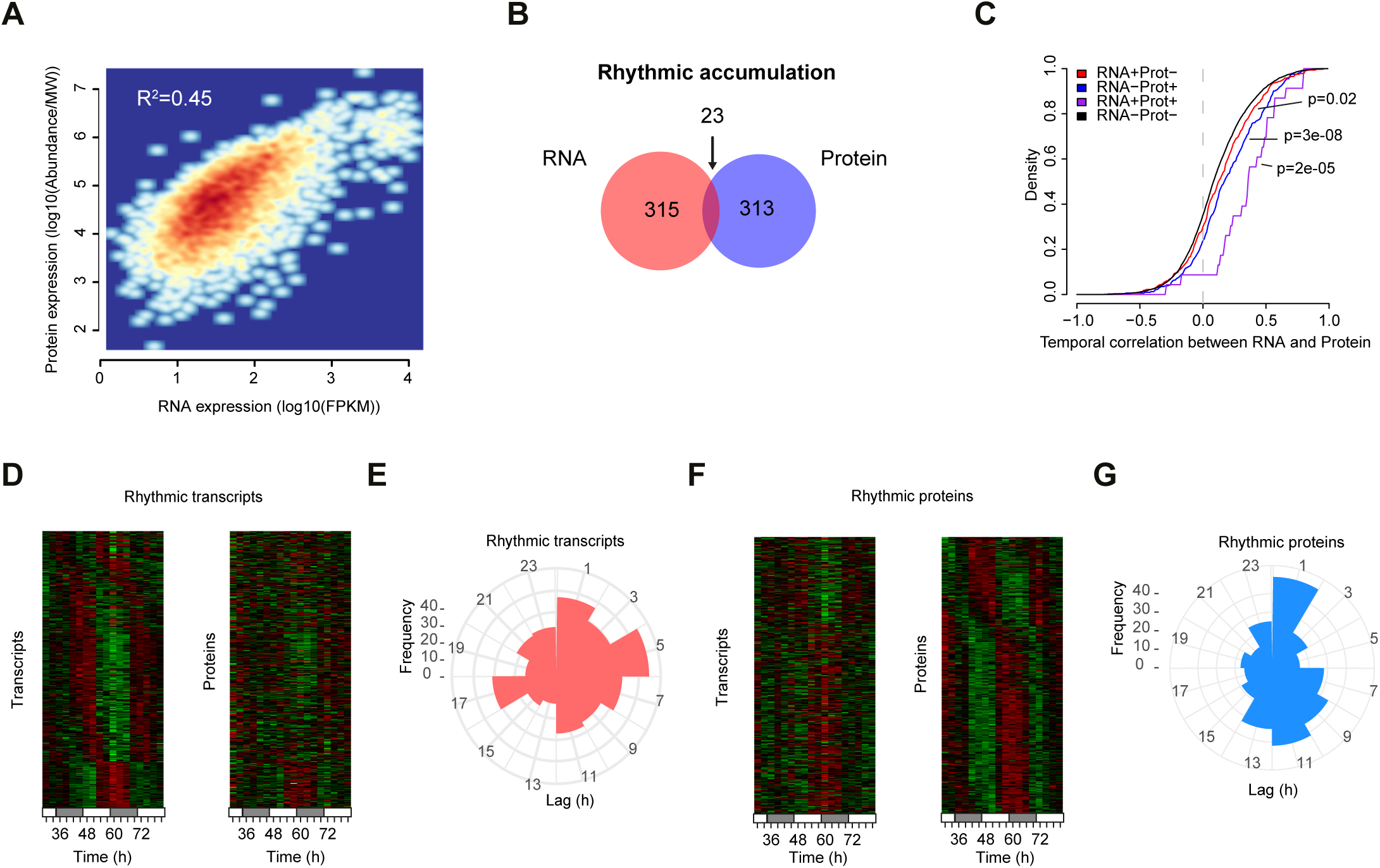
Integration of transcriptomics and proteomics data. (A) Scatter plot of RNA and protein expression. (B) Venn diagram showing the overlap of rhythmic RNA and proteins. (C) Cumulative distributions for Pearson correlation coefficients for the indicated groups of RNA-protein pairs. RNA-Prot-, neither circadian; RNA+Prot+, both circadian; RNA-Prot+, RNA not circadian and protein circadian; RNA+Prot-, RNA circadian and protein not circadian (Wilcoxon sum rank test). (D and F) Heatmap representations of transcripts and protein accumulations for the RNA-protein pairs with rhythmic transcripts (D) or proteins (F). (E and G) Distribution of phase difference between RNA-protein pairs for those with rhythmic transcripts (E) or proteins (G).

We reached similar conclusions when looking at the temporal cross-correlation between transcripts and proteins (Figure S3A). In particular, we found that the set of RNA-protein pairs with rhythmic transcripts had a wide distribution of phase lags between RNA and protein, suggesting that the phase of transcripts is a poor predictor of protein accumulation (Figure 3D-E). In contrast, rhythmic proteins tended to be expressed in phase with their transcripts, or with a phase delay of about 12 h (Figure 3F-G). This phase delay is likely related to the fact that most transcripts are expressed around CT12 (Figure 1D), while most proteins peak around CT0 (Figure 2C). We further explored the global temporal patterns of our transcriptome and proteome datasets using principal component analysis (PCA). Interestingly, ordered temporal transitions between four different transcriptional states were mostly captured by the first two PCA components (Figure S3B). When considering matching protein profiles, we observed similar clustering of time points, but this time with three states showing a cyclic relationship (Figure S3C). Together, these results highlight the combination of transcriptional and post-transcriptional mechanisms in the regulation of the circadian proteome.

### Central carbon metabolism and amino acid metabolism exhibit daily oscillations

Our analysis of the circadian transcriptome and proteome highlighted metabolism as a key cellular function regulated in a circadian manner. We therefore considered that it would be informative to determine which metabolites might exhibit 24-hour oscillation. First, we performed untargeted liquid chromatography-mass spectrometry (LC-MS) and detected 1339 features, among which 466 were rhythmic (Figure 4A and Figure S4A-B; JTK-Cycle, adjusted p-value < 0.05; FDR = 14.5%). In keeping with our gene expression data, we found that rhythmic features clustered into two phases (Figure 4B-C), indicating a high degree of temporal organisation. Using tandem MS (MS2), we were able to identify 54 metabolites and performed pathway enrichment analysis (Figure 4D, Figure S4C and Table S1; Metabolite Set Enrichment Analysis, raw p-value < 0.05) (23). Importantly, central carbon metabolism and pathways associated with amino acid metabolism were enriched among rhythmic metabolites. Given this, we further investigated the temporal regulation of central carbon metabolism by performing targeted LC-MS analysis of glycolysis, pentose phosphate pathway and citric acid cycle metabolites. This revealed that key metabolites including ATP, glutathione and citric acid cycle intermediates exhibited rhythmic accumulation over time (Figure 4E-F and Table S2).

**Figure 4.**
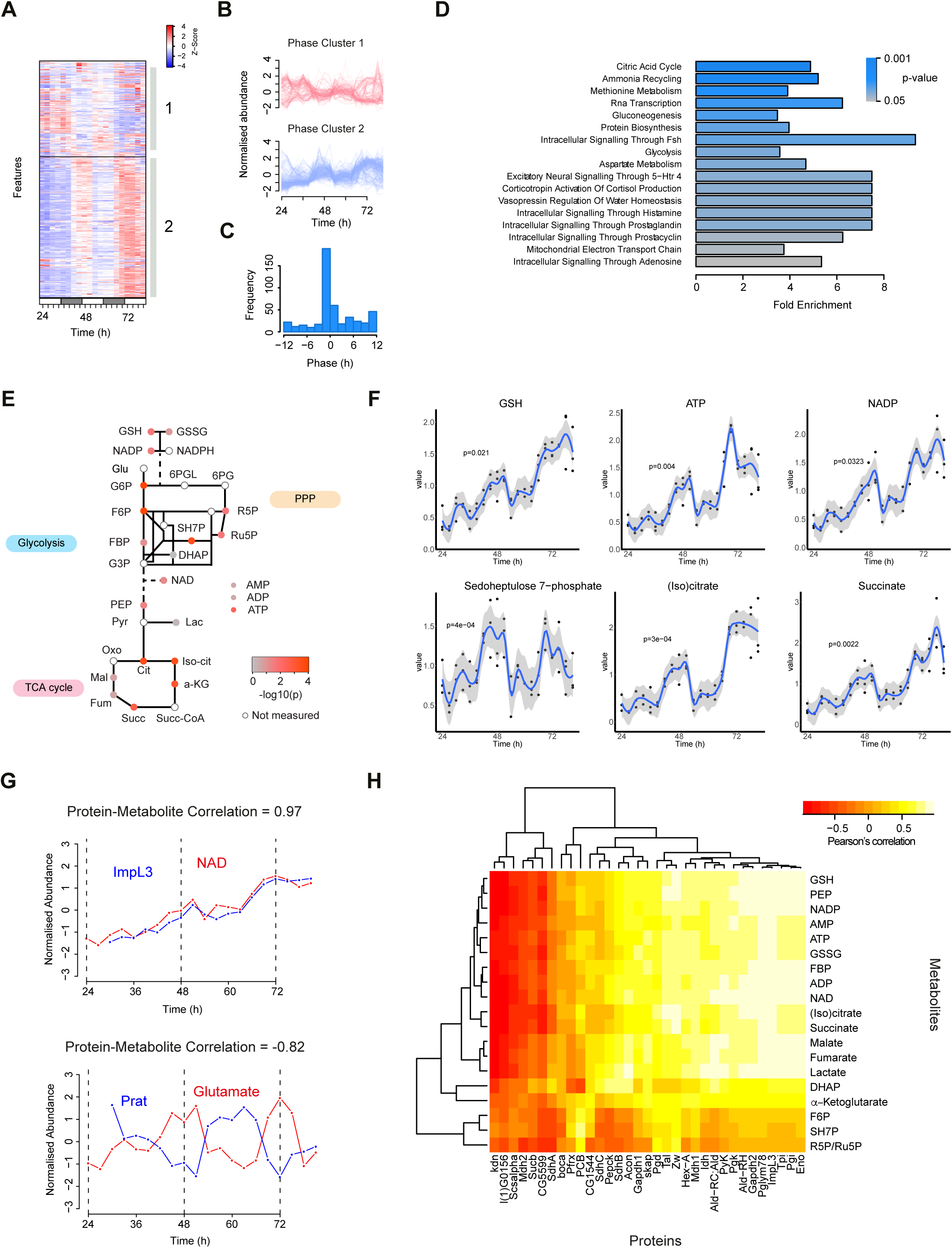
The circadian metabolome of S2 cells. (A) Heatmap representation of the 466 LC-MS features detected as rhythmic (JTK-Cycle; P<0.05, FDR=14.5%). (B) Individual traces of rhythmic features from cluster 1 and 2 from (A). (C) Distribution of phases from (A). (D) Metabolite Set Enrichment Analysis of the 54 identified rhythmic metabolites. (E) Targeted LC-MS analysis of metabolites from glycolysis, pentose phosphate pathway (PPP) and tricarboxylic acid (TCA) cycle. The colour of each node represents the p-value for daily rhythmicity (JTK Cycle). See Table S2 for list of abbreviations. (F) Selected metabolite temporal profiles are shown together with their associated p-value. (G) Two examples of protein-metabolite pairs are shown with their respective Pearson correlation coefficients. (H) Heatmap showing the Pearson ‘s correlation between each profile from targeted metabolomics (carbon metabolism) and proteomics data.

In order to infer relationships between rhythmic protein expression and metabolic oscillations, we integrated our proteomics dataset with the tandem MS-annotated metabolite data, and performed correlation analyses. When considering all possible protein to metabolite associations, the distribution of correlation coefficients displayed a non-uniform profile with an overrepresentation of highly correlated and anti-correlated profiles (Figure S4D). When only the best match between each metabolites and associated proteins was kept, the effect was much more pronounced, since most protein-metabolite pairs had an absolute correlation coefficient greater than 0.5 (Figure S4E). For example, lactate dehydrogenase (*ImpL3*) and NAD displayed strongly correlated profiles (Figure 4G; Pearson ‘s correlation = 0.97), while glutamate accumulation was anti-correlated to the levels of the phosphoribosylamidotransferase (Prat), an essential enzyme in the pathway for *de novo* purine synthesis (Pearson ‘s correlation = -0.82). Using a similar strategy, we analysed the correlation patterns between proteins and targeted metabolites from central carbon metabolism (Figure 4H). This revealed that most protein-metabolite pairs were correlated, especially those found in the same sub-pathway. For example, proteins and metabolites of the pentose phosphate pathway formed a small cluster that exhibited correlated profiles, as exemplified by sedoheptulose 7-phosphate and transaldolase (Taldo) (Figure S4F). Together, these results imply that there is a robust link between protein rhythmicity and downstream metabolite oscillations in key metabolic pathways.

## Discussion

Current models of circadian oscillations rely on transcription-translation feedback loops, in which the core transcription factors CLK and CYC (CLOCK and BMAL1 in mammals) occupy a central role. Here we show that *Drosophila* S2 cells do not express the core components of the canonical transcriptional loops, but nevertheless exhibit genome-wide oscillations at the level of transcripts, proteins and metabolites. Therefore, this novel system extends the notion of circadian oscillations beyond the classical framework and thus opens new avenues to explore the fundamental underpinnings of circadian oscillations in a genetically-tractable system.

Studies in mammalian system have suggested that canonical circadian genes may not be necessary for the generation of circadian oscillations. Indeed, human red blood cells, which lack a nucleus and thus transcription, display self-sustained redox oscillations on the circadian time scale (14). Moreover, the study of circadian oscillations during embryonic differentiation has led to the surprising observation that circadian rhythmicity precedes the normal expression of clock genes, for example circadian glucose uptake in mouse undifferentiated stem cells (11) and rhythmic gene expression in human embryonic stem cell-derived cardiomyocytes (12).

Together, these results indicate that the canonical circadian network may not be primarily involved in time keeping itself, but rather involved in the coordination of cellular oscillation (Figure 5). The fact that CLK/CYC are transcription factors with a PAS domain, which allow molecular sensing of the intracellular environment, is consistent with a role as an important cog between metabolic and gene expression programs. In this interpretation, *Drosophila* S2 cells are thus a novel model of cellular time keeping that encompasses the metabolic and gene expression components, but not the circadian network. Understanding the relative role of each part in this model will surely enhance greatly our understanding of fundamental properties of circadian oscillations.

**Figure 5.**
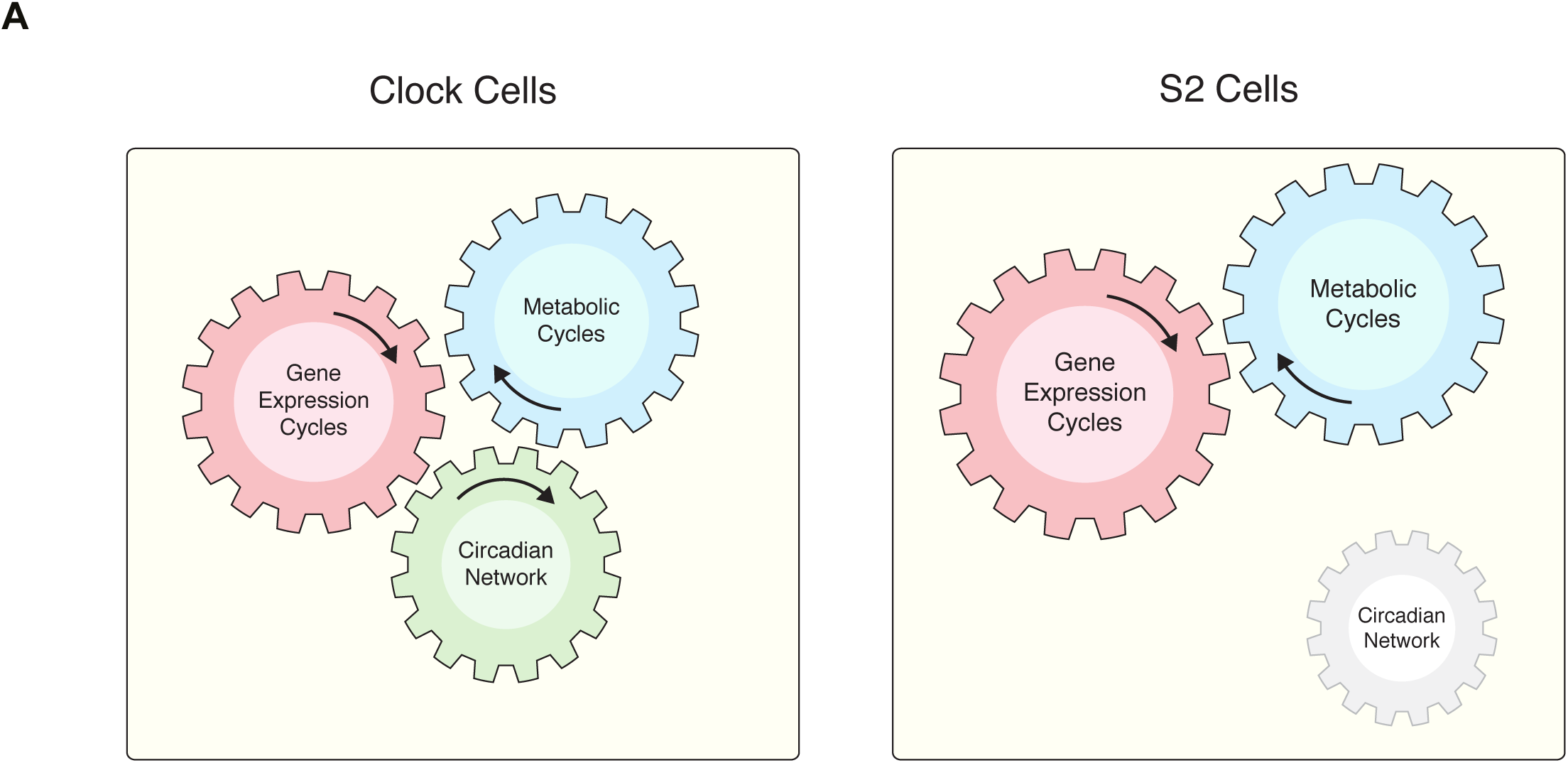
Coupled gene expression and metabolic cycles in *Drosophila* S2 cells. (A) Schematic showing the difference between a clock cell where the circadian network is active and *Drosophila* S2 cells, which exhibit gene expression and metabolic cycles in its absence.

In unicellular organisms such as yeast and cyanobacteria, gene expression cycles coupled to metabolic oscillations have been described with a period of oscillation in the hour range (24, 25). These short-period metabolic oscillations are considered by some as a prototype of circadian oscillations (26). Of particular interest, a recent study showed that such metabolic oscillations constitute a single-cell phenomenon, since they have been reported in single yeast cells recorded in a microfluidic device (27). Importantly, such metabolic oscillations are not driven by the cell cycle, but rather they gate the progression of the latter. Accordingly, we did not observe a significant variation in the proportion of cell cycle phases over time, indicating that a similar relationship between circadian oscillations and cell cycle may take place in S2 cells.

The finding that energy metabolism and protein translation are regulated in a circadian fashion in *Drosophila* S2 cells suggests that the two processes are intimately linked, even in absence of a functional circadian network. Therefore, we put forward the hypothesis that gene expression cycles are coupled to metabolic oscillations in order to synchronise energetically costly protein biosynthesis with energy metabolism. In this context, a recent study has proposed that rhythmic gene expression optimizes the metabolic cost of global gene expression and that highly expressed genes have been selected to be downregulated in a cyclic manner for energy conservation (28). Therefore, bioenergetic constraints are likely to play an important role in *Drosophila* cells and may be an important driver of circadian oscillations in this, and other, eukaryotic models.

## Materials and Methods

### Cell culture

Drosophila S2 cells were purchased from Thermo Fisher Scientific and were grown in Schneider ‘s Drosophila Media (Thermo Fisher Scientific), supplemented with 10% Heat-inactivated FBS, 1% Pen-Strep and 1/500 MycoZap Plus-CL. For circadian entrainment protocol, S2 cells were subjected to temperature cycles (12 h at 23°C, 12 h at 28°C) for at least one week, with media changes occurring every 3-4 days at the transition between 23 to 28°C. The last medium change was performed at t = 0 h and cells were plated into 6-well plates. Cells were kept at 25°C for the remaining of the experiment, with sample collection occurring at 3 h intervals between 24 h and 81 h.

### RNA isolation and RNA-Sequencing

At the time points indicated in the main text, cells were lysed in triplicate in TRI-Reagent, flash-frozen and stored at -80°C until extraction. Extraction and purification were performed with the Direct-zol RNA MiniPrep kit (Zymo Research). RNA-seq libraries were prepared as described previously (29). Sequencing using a HiSeq platform with single-end 50 bp reads and subsequent quality filtering of reads was performed according to manufacturer ‘s instructions (Illumina).

### RNA-Seq data analysis

Sequencing reads were aligned to the UCSC Drosophila reference genome (dm3) using TopHat v2.1.0 (30). Reads were assembled into transcripts using RefSeq genes as reference and their abundance estimated using Cufflinks/Cuffmerge/Cuffquant/Cuffnorm v2.2.1 (31). To estimate the threshold to define the set of expressed transcripts, we modeled the distribution of transcript mRNA expression using a Gaussian mixture model using the R package “mixtools". We chose a threshold of 0.9 FPKM, which corresponds to the 0.95 percentile of the distribution of lowly expressed transcripts, to define a set of 6944 expressed transcripts.

Temporal profiles were linearly detrended and the JTK-Cycle algorithm was used to detect rhythmic transcripts using the following parameters: minimal period = 21, maximal period = 27, adjusted p-value = 0.05. As validation, two alternative algorithms, RAIN (period = 24, period.delta = 3, method = longitudinal, and p-value = 0.01) and ARSER (minimal period = 21, maximal period = 27, default period = 24 and p-value = 0.01) were used to detect circadian transcripts. GO analyses were performed using the Gene List Analysis Tool from the PANTHER database using all *D. Melanogaster* genes as reference set (32).

### Quantitative PCRss

Triplicate RNA samples were reverse-transcribed into cDNA using the High Capacity cDNA Reverse Transcription Kit (Life Technologies), following the manufacturer ‘s instructions, using 0.3-1µg total RNA per reaction. The resulting cDNAs were used in duplicate 7 µL PCR reactions, set up as follows: 3.5 µL TaqMan Gene Expression Master Mix (Life Technologies), 0.35 µL validated Taqman Gene Expression Assay (Life Technologies), 1.15 µL nuclease-free water and 2 µL cDNA. Real-time PCR was performed with an ABI 7900HT (Applied Biosystems) system. The following Taqman Gene Expression Assays were used: Eno (Dm01844953), Gapdh2 (Dm01843776), ImpL3 (Dm01841229) and Act5C (Dm02361909). The relative levels of each mRNA were calculated by the 2^-**Δ**Ct^ method and normalized to the corresponding Act5C levels.

### Flow cytometry

At the time points mentioned in the main text, cells were washed twice in PBS and resuspended in ice-cold 70% Ethanol. Cells were kept at 4°C until staining. For staining DNA, cells were first washed twice with PBS, resuspended in DAPI solution (1 mg/mL in PBS + 0.1% Triton) and kept overnight at 4°C. Flow cytometry was performed on a LSRFortessa™ cell analyzer (BD Biosciences) using standard methods.

### Proteomics sample preparation

At the time points indicated in the main text, cells were spun down and the pellets were flash-frozen and stored at -80°C until protein extraction. To extract protein, pellets were lysed on ice with 500 µL of Lysis Buffer (100 mM Triethylammonium bicarbonate (TEAB), 1% SDS, 1% NP-40, 10 mM diethylenetriaminepentaacetic acid (DTPA),1/100 Halt protease inhibitors (Thermo Fisher Scientific)). Cells were vortexed and incubated for 30 min on ice. Samples were sonicated using a Bioruptor Standard (Diagenode) for 5 min (30 s On, 30 s Off) on medium power. Samples were spun at max speed at 4°C for 10min to remove debris and transferred to fresh tubes. BCA assay (Thermo Fisher Scientific) was used to quantify protein levels for tandem-mass tag (TMT) labelling (Thermo Fisher Scientific).

TMT labelling was performed according to manufacturer ‘s instructions. 200 µg per condition was transferred into a new tube and the volume was adjusted to 200 µL with 100 mM TEAB. 10 µL of 200 mM TCEP was added to each sample to reduce cysteine residues and samples were incubated at 55°C for 1 h. To alkylate cysteines, 10 µL of 375 mM iodoacetamide was added to each sample and samples were incubated for 30 min protected from light at room temperature. Samples were split in two and acetone precipitation was performed by adding 6 volumes (∼600 µL) of pre-chilled (-20°C) acetone. The precipitation was allowed to proceed overnight at -20°C. The samples were centrifuged at 8000 × *g* for 10 min at 4°C, before decanting the acetone.

Acetone-precipitated (or lyophilized) protein pellets were resuspended with 100 µL of 100 mM TEAB. 2.5 µg of trypsin per 100 µg of protein was added to the proteins for proteolytic digestion. Samples were incubated overnight at 37°C to complete the digestion. TMT Label Reagents were resuspended in anhydrous acetonitrile and 0.4 mg of each label was added to the corresponding peptide sample. The reaction was allowed to proceed for 1 h at room temperature. 8 µL of 5% hydroxylamine was added to each sample and incubated for 15 min to quench the labelling reaction. Samples were combined in a new microcentrifuge tube at equal amounts and store at -80°C until mass spectrometry analyses.

### Proteomics mass spectrometry

TMT-labelled tryptic peptides were subjected to HpRP-HPLC fractionation using a Dionex Ultimate 3000 powered by an ICS-3000 SP pump with an Agilent ZORBAX Extend-C18 column (4.6 mm × 250 mm, 5 µm particle size). Mobile phases (H_2_0, 0.1% NH_4_OH or MeCN, 0.1% NH_4_OH) were adjusted to pH10.5 with the addition of formic acid and peptides were resolved using a linear 40 min 0.1–40 % MeCN gradient over 40 min at a 400 µL/min flow rate and a column temperature of 15°C. Eluting peptides were collected in 15 s fractions. 120 fractions covering the peptide-rich region were re-combined to give 12 samples for analysis. To preserve orthogonality, fractions were combined across the gradient. Re-combined fractions were dried down using an Eppendorf Concentrator (Eppendorf, UK) and resuspended in 15 µL MS solvent (3% MeCN, 0.1% TFA).

Data for TMT labelled samples were generated using an Orbitrap Fusion Tribrid Lumos mass spectrometer (Thermo Scientific). Peptides were fractionated using an RSLCnano 3000 (Thermo Scientific) with solvent A comprising 0.1% formic acid and solvent B comprising 80% MeCN, 20% H_2_O, 0.1% formic acid. Peptides were loaded onto a 75 cm Acclaim PepMap C18 column (Thermo Scientific) and eluted using a gradient rising from 7 to 37 % solvent B by 180 min at a flow rate of 250 nL/min. MS data were acquired in the Orbitrap at 120,000 fwhm between 380–1500 m/z. Spectra were acquired in profile with AGC 2 × 10^5^. Ions with a charge state between 2+ and 7+ were isolated for fragmentation using the quadrupole with a 0.7 m/*z* isolation window. CID fragmentation was performed at 35% collision energy with fragments detected in the ion trap between 400–1200 m/*z*. AGC was set to 1 × 10^4^ and MS2 spectra were acquired in centroid mode. TMT reporter ions were isolated for quantitation in MS3 using synchronous precursor selection. Ten fragment ions were selected for MS3 using HCD at 65% collision energy. Fragments were scanned in the Orbitrap at 60,000 fwhm between 120–500 m/*z* with AGC set to 1 × 10^5^. MS3 spectra were acquired in profile mode.

### Proteomics data analysis

MaxQuant v1.5.5.1 (33) was used to process the raw TMT proteomics data using the following parameters: Fixed modifications = carbamidomethylation, FDR for protein and peptide identification = 0.01, sequence database=UniprotKB proteome for *Drosophila melanogaster* (downloaded on 13 January 2017), variable modifications= oxidation of methionine, protein N-terminal acetylation. TMT 10plex data were normalised using linear regression in logarithmic space to normalise for global abundance variations between each TMT channel. For the quantification TMT sets, samples were TMT-labelled as shown in Figure 2A (set1: pooled sample, CT30, CT33, CT36, CT39, CT42, CT45, CT48, CT51, CT54; set2: pooled sample, CT57, CT60, CT63, CT66, CT69, CT72, CT75, CT78, CT81). For each protein, the quantification TMT set1 and set2 were assembled together after taking the log ratio between each channel and the reference channel (pooled sample of all time points). For the validation TMT sets, samples were TMT-labelled as shown in Figure S2C (set1: CT24, CT30, CT36, CT42, CT48, CT54, CT60, CT66, CT72, CT78; set2: CT27, CT33, CT39, CT45, CT51, CT57, CT63, CT69, CT75, CT81). For each protein, the validation TMT set1 and set2 were assembled together after taking the log ratio between each channel and the mean across all channels for this protein (mean centering). Protein temporal profiles were linearly detrended and the JTK-Cycle algorithm (16) together with the RAIN and ARSER methods for validation were used to detect rhythmic transcripts with same parameters used for the RNA-Seq analysis. GO analyses were performed using the Gene List Analysis Tool from the PANTHER database using all *D. Melanogaster* genes as reference set (32). In order to integrate transcriptomics and proteomics data, we used the Uniprot ID converter tools to generate a mapping between Uniprot accession numbers and RefSeq annotations. We performed a protein-centric integration, where each protein was associated with at most one transcript. If there were more than one transcript, the most rhythmic mRNA transcript was kept. Using this strategy, we were able to assemble 4658 protein-RNA pairs from 4758 proteins and 6944 transcripts. For each RNA-protein pair, the Pearson correlation coefficient was computed using the respective temporal profiles from CT30 to CT81.

### Metabolite sample preparation

At the time points indicated in the main text, triplicate cell samples were spun down and pellets were washed with room-temperature PBS. Cells were resuspended in 1 mL of methanol:water (80:20) at - 75°C for quenching and samples were stored at -80°C. To extract metabolites, samples were thawed on ice and then vortexed for 2 min at room temperature. Samples were sonicated for 5 min in the cold room (30 s ON, 30 s OFF, medium power). Samples were centrifuged the mixture for 10 min at 10,000 rpm at 4°C and the supernatants were transferred to fresh 1.5 mL Eppendorf tubes. The extraction was repeated a second time with 500 µL methanol:water (80:20). For the third extraction, 500 µL methanol:water (80:20) supplemented with ^13^C_5_ ^15^N_1_ -valine was used downstream quality control. The three extractions were pooled and lyophilized to dryness. Samples were resuspended in 350 µL of chloroform:methanol:water (1:3:3 v/v) and the polar (upper) phase was collected for analysis. Quality control samples were prepared by pooling equal volumes from all samples included in this study.

### Metabolomics mass spectrometrys

LC-MS method was adapted from a published protocol (34). Samples were injected onto a Dionex UltiMate LC system (Thermo Scientific) with a ZIC-pHILIC (150 mm x 4.6 mm, 5 µm particle) column (Merck Sequant). A 15 min elution gradient of 80% to 20% Solvent B was used, followed by a 5 min wash of 5% Solvent B and 5 min re-equilibration, where Solvent B was acetonitrile (Optima HPLC grade, Sigma Aldrich) and Solvent A was 20 mM ammonium carbonate in water (Optima HPLC grade, Sigma Aldrich). Other parameters were as follows: flow rate 300 µL/min; column temperature 25°C; injection volume 10 µL; autosampler temperature 4°C. MS was performed with positive/negative polarity switching using an Q Exactive Orbitrap (Thermo Scientific) with a HESI II probe. MS parameters were as follows: spray voltage 3.5 kV and 3.2 kV for positive and negative modes, respectively; probe temperature 320°C; sheath and auxiliary gases were 30 and 5 arbitrary units, respectively; full scan range: 70 to 1050 m/*z* with settings of AGC target and resolution as Balanced and High (3 × 10^6^ and 70,000) respectively. Data was recorded using Xcalibur 3.0.63 software (Thermo Scientific). Mass calibration was performed for both ESI polarities before analysis using the standard Thermo Scientific Calmix solution. To enhance calibration stability, lock-mass correction was also applied to each analytical run using ubiquitous low-mass contaminants. Parallel reaction monitoring (PRM) acquisition parameters: resolution 17,500; collision energies were set individually in HCD (high-energy collisional dissociation) mode.

### Metabolomics data analysis

Qualitative analyses were performed using Xcalibur Qual Browser (Thermo Fisher Scientifc) and mzCloud (HighChem). Untargeted metabolomics data analyses were performed with Progenesis QI (Nonlinear Dynamics) using the following parameters: feature detection = high resolution, peak processing = centroided data with resolution at 70000 (FWHM). In positive mode, the following adducts were used: M+NH_4_, M+H, M+Na and M+2H. In negative mode, the following adducts were used: M-H, M+Na-H, M-2H. Normalisation was performed using the log-ratio method over all features. Features having a coefficient of variation (CV) lower than 30% among quality control samples were selected for downstream analyses (n = 722 and 616 for positive and negative mode, respectively). PCA of all samples (including features with CV < 30% from positive and negative modes) shows very good clustering, indicating system stability, performance, and reproducibility (Figure S4A). Similar conclusions were reached using correlation analysis (Figure S4B). Features in the retention time window between 19.15 and 19.35 min were excluded from subsequent analyses, due to artefactual profiles in this time window. Temporal profiles were linearly detrended and the JTK-Cycle algorithm was used to detect circadian rhythmicity using the following parameters: minimal period = 21, maximal period = 27, adjusted p-value = 0.05, number of replicates = 2-3. From the 466 rhythmic features, 145 with at least one hit in spectral databases were selected for MS2 annotation. Out of these, we were able to annotate 70 features with MS2 data (Table S1), which correspond to 54 metabolites. Metabolic pathway enrichment analysis was performed using Metabolite Set Enrichment Analysis (MSEA) (23). For targeted LC-MS data analysis, a set of 20 metabolites (Table S2) was chosen from carbon metabolism and redox pathways. Retention time and MS/MS spectra from samples were compared to metabolite standards to validate identification. Quantification was performed manually using TraceFinder v4.1 (Thermo Fisher Scientific). Normalisation across samples was performed using the normalisation ratio calculated with Progenesis QI. In order to integrate metabolomics and proteomics datasets, we used the Kyoto Encyclopedia of Genes and Genomes (KEGG) annotation. Briefly, Uniprot accession numbers were annotated with Enzyme Commission (EC) numbers, which were used to fetch all interacting metabolites in the KEGG database. Each metabolite was annotated with all possible proteins based on the described annotation and correlation analysis was performed between metabolite-protein pairs.

### Data deposition

The RNA-seq data set produced in this study has been deposited in the Gene Expression Omnibus (accession number GSE102495). The mass spectrometry proteomics data have been deposited to the ProteomeXchange Consortium via the PRIDE (35) partner repository with the dataset identifier PXD007669.

## Acknowledgments

ABR acknowledges funding from the Wellcome Trust (100333/Z/12/Z), the European Research Council (ERC Starting Grant No. 281348, MetaCLOCK), EMBO Young Investigators Programme, the Lister Institute of Preventive Medicine. ABR is supported by The Francis Crick Institute, which receives its core funding from Cancer Research UK (FC001534), the UK Medical Research Council (FC001534), and the Wellcome Trust (FC001534). GR was supported by an Advanced SNSF Postdoctoral Mobility Fellowship and an EMBO Long-Term Fellowship.

## Supplemental Information

**Figure S1.**
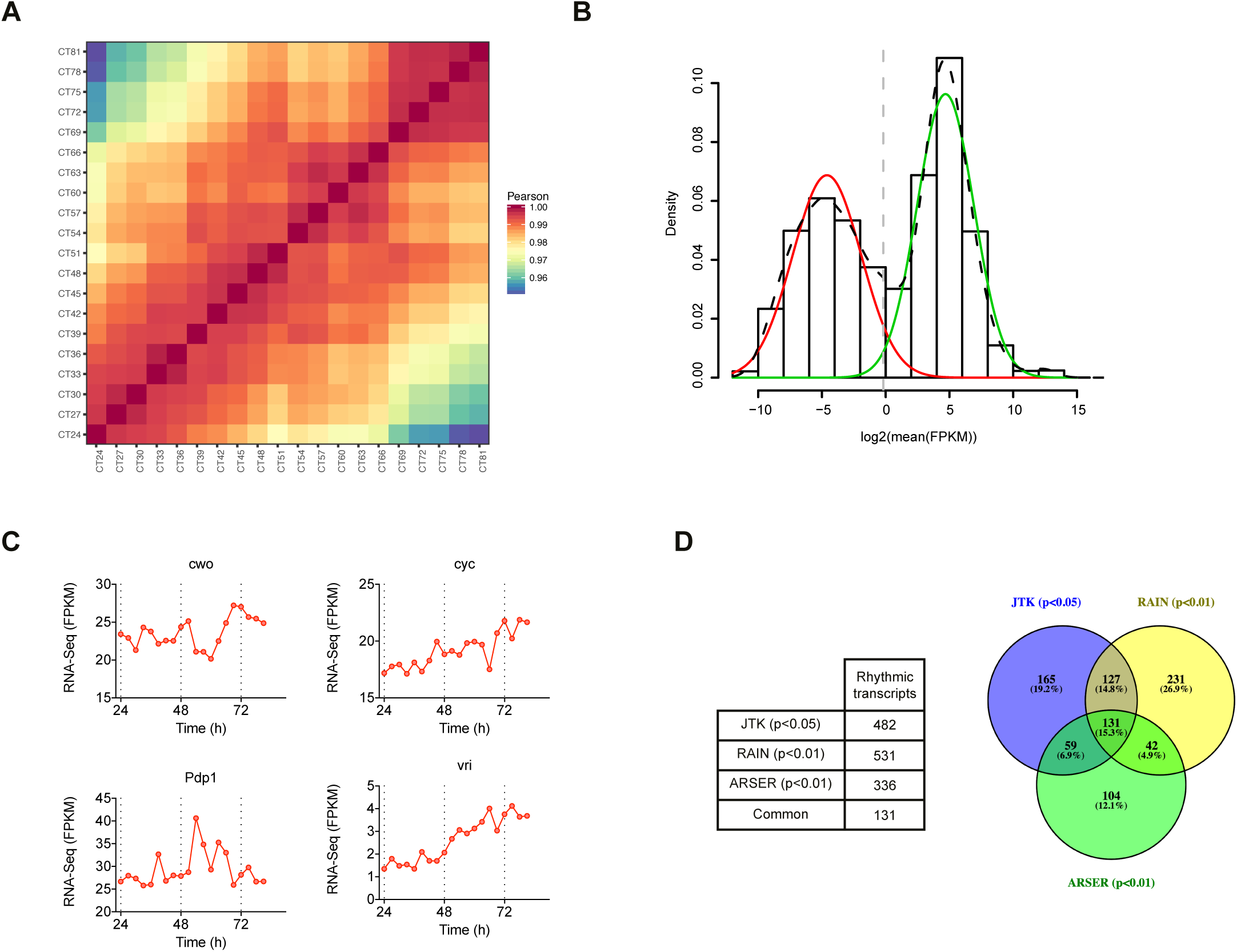
(A) Heatmap showing the Pearson ‘s correlation coefficient between each pair of time points from the RNA-Seq time course. (B) Histogram showing the distribution of mean expression levels measured by RNA-Seq. The distribution was modelled with a mixture model to define the sets of lowly (red) and highly (green) expressed transcripts. The dashed vertical line represents the cut-off chosen to define the set of expressed transcripts. FPKM, Fragments Per Kilobase of transcript per Million mapped reads. (C) Validation of the JTK-Cycle algorithm with two other methods to detect rhythmic transcripts. A table giving the number of rhythmic transcripts is given (left), together with a Venn diagram showing the overlap between methods (right). (D) mRNA expression of four canonical circadian expressed in S2 cells.

**Figure S2.**
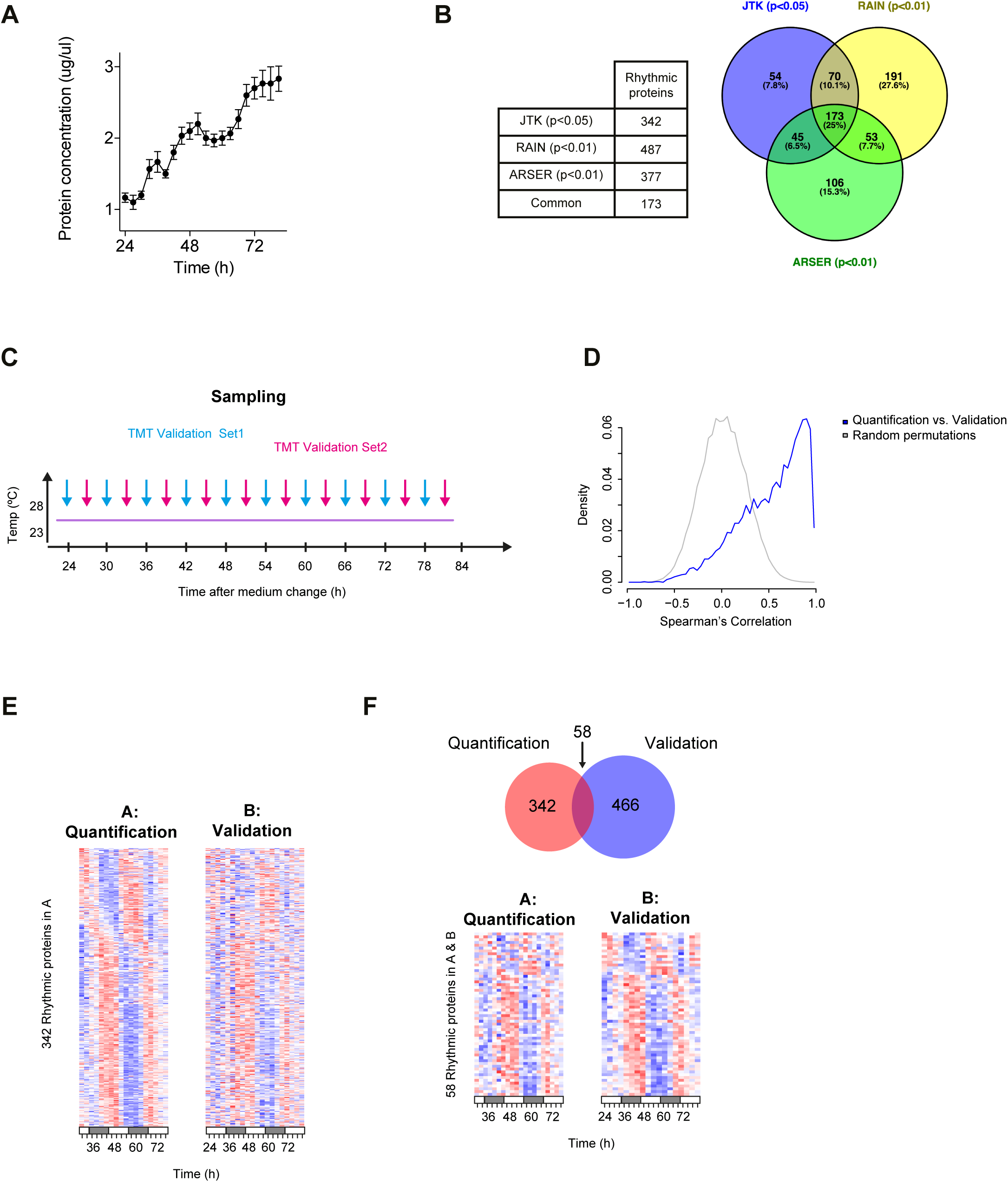
(A) Validation of the JTK-Cycle algorithm with two other methods to detect rhythmic transcripts. A table giving the number of rhythmic transcripts is given (left), together with a Venn diagram showing the overlap between methods (right). (B) The protein concentration in cell extracts was measured using BCA method (mean ± SEM, n = 2-3). (C) Cartoon representing the labelling scheme for the validation protocol. Samples were labelled using interspersed time points (Set 1: CT24, CT30, CT36,…; Set 2: CT27, CT33, CT39,…) and the two 10plex TMTs were then combined to produce 20 time point quantification profiles. (D) Distributions of Spearman ‘s correlation coefficient between the quantification and validation sets of quantitative proteomics data. Random permutations of the data are shown as control (n = 100). (E) Heatmap representation of the proteins detected as rhythmic in the quantification set (342 proteins) for the quantification set (left) and validation set (right). (F) Heatmap representation of the proteins detected as rhythmic in both the quantification and validation sets (58 proteins) for the quantification set (left) and validation set (right).

**Figure S3.**
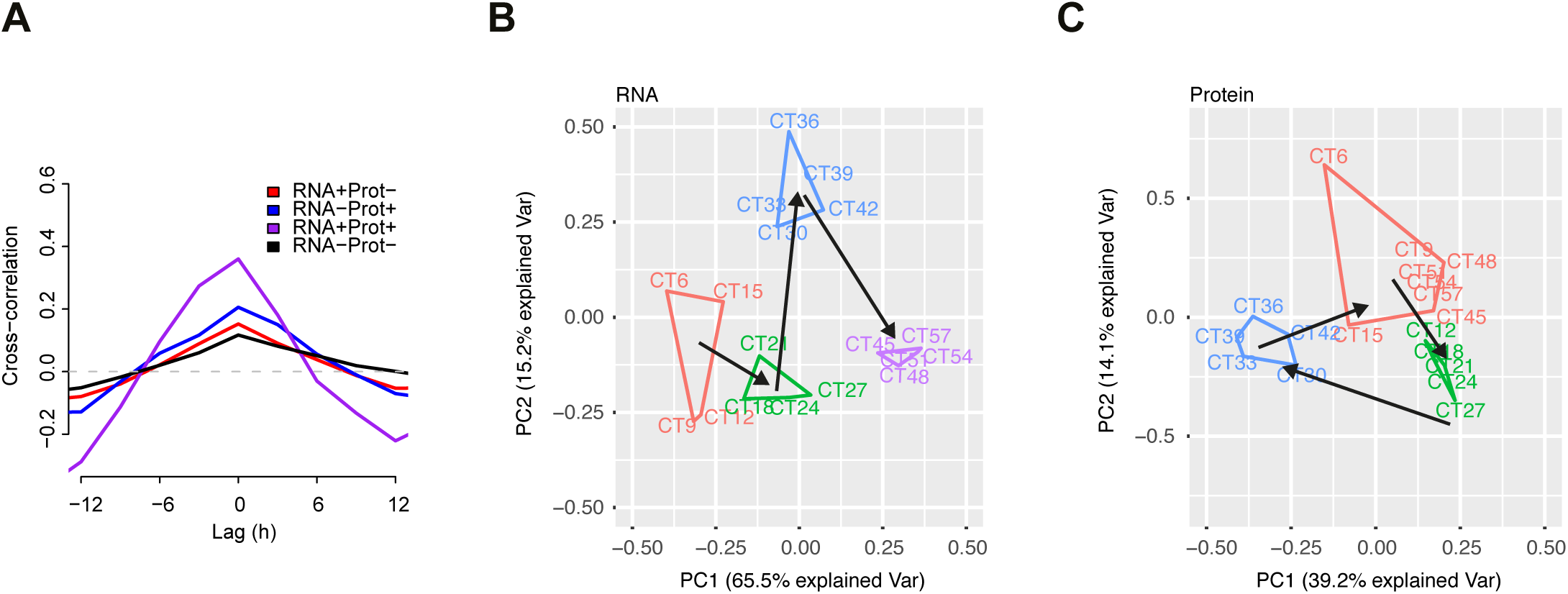
(A) Temporal cross-correlation for RNA-proteins pairs as described in Figure 3. (B-C) PCA plot of RNA and protein data for all RNA-proteins pairs. Clustering of time points is performed with Partitioning Around Medoids (PAM) method.

**Figure S4.**
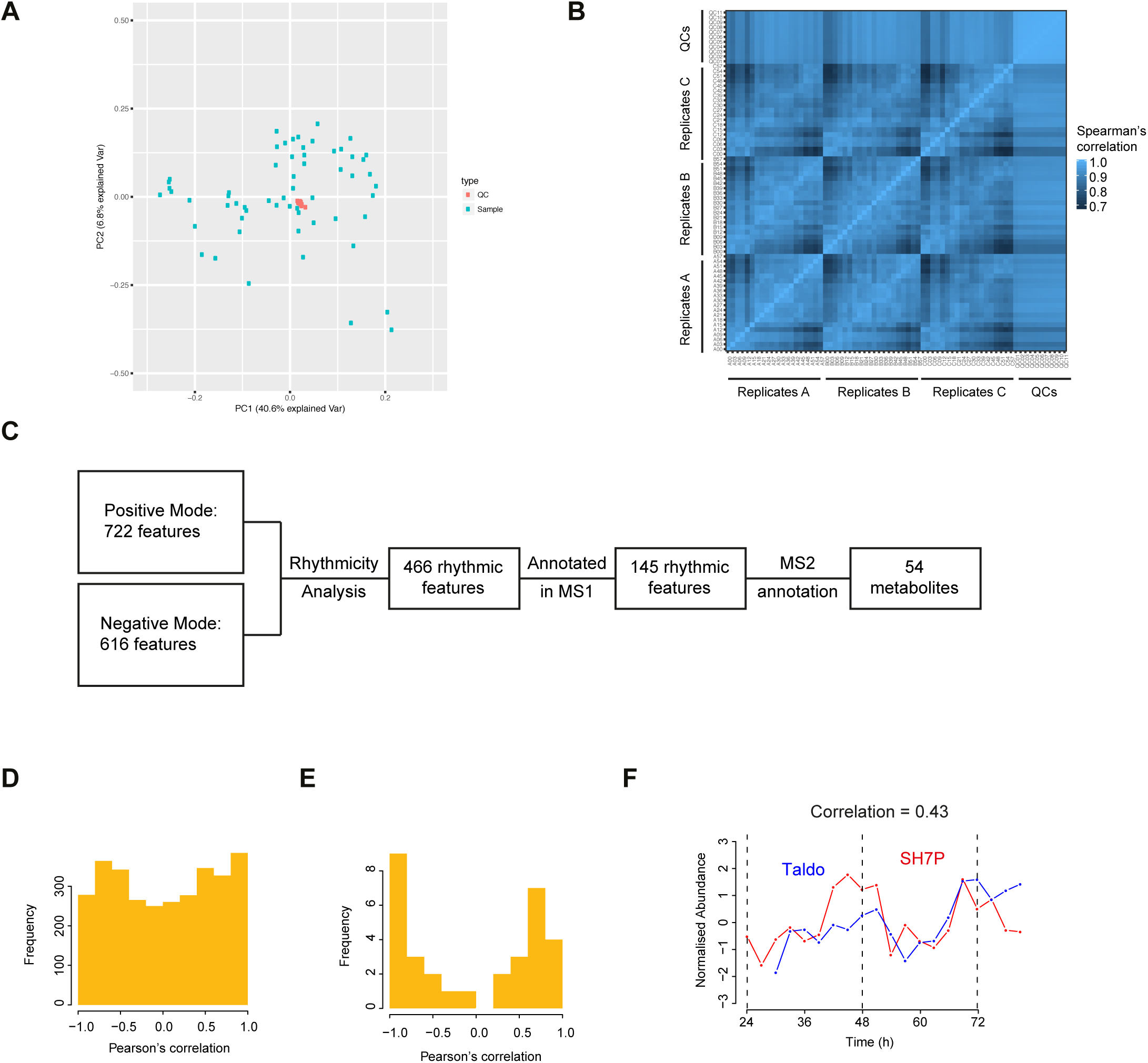
(A) PCA plot of the 59 samples and 11 quality control (QC) samples analysed in this study. (B) Heatmap showing the Spearman ‘s correlation between all samples. (C) Workflow of untargeted metabolomics data analysis. (D) Distribution of Pearson ‘s correlation coefficients for all possible protein-metabolite pairs. (E) Distribution of Pearson ‘s correlation coefficients for the best protein-metabolite pairs. For each metabolite, the correlation coefficients for all possible interacting proteins were computed and the protein with the highest absolute correlation coefficient was retained. (F) An examples of correlated protein-metabolites are shown with Pearson correlation coefficient (top). SH7P, sedoheptulose 7-phosphate; Taldo, Transaldolase.

**Table S1.** Table of the 70 MS2-annotated features with m/z, retention time and fragments used for annotation.

| m/z | Retention time (min) | Charge | Polarity | MS2 Annotation | CAS | KEGG | Fragments |
| --- | --- | --- | --- | --- | --- | --- | --- |
| 132.07 | 12.17 | 1 | Positive | 3-Hydroxy-L-proline | 4298-08-2 | C19706 | 90.06, 132.08 |
| 136.05 | 9.78 | 1 | Positive | Adenine | 73-24-5 | C00147 | 119.04, 92.02, 136.06 |
| 508.00 | 12.85 | 1 | Positive | Adenosine triphosphate | 56-65-5 | C00002 | 136.06, 410.03, 428.04 |
| 118.09 | 10.80 | 1 | Positive | Betaine | 107-43-7 | C00719 | 118.09 |
| 332.14 | 8.15 | 1 | Positive | Ciprofloxacin | 85721-33-1 | C05349 | 231.06, 332.14 |
| 489.11 | 12.79 | 1 | Positive | Citicoline | 987-78-0 | C00307 | 264.04, 184.07 |
| 223.07 | 13.56 | 1 | Positive | Cystathionine | 535-34-2 | C00542 | 134.08, 223.08 |
| 244.09 | 11.22 | 1 | Positive | Cytidine | 65-46-3 | C00475 | 95.02, 112.05 |
| 324.06 | 13.27 | 1 | Positive | cytidine 5-monophosphate | 63-37-6 | C00055 | 112.05, 324.06 |
| 112.05 | 13.27 | 1 | Positive | Cytosine | 71-30-7 | C00380 | 95.02, 112.05 |
| 112.05 | 11.22 | 1 | Positive | Cytosine | 71-30-7 | C00380 | 95.02, 112.05 |
| 148.06 | 12.68 | 1 | Positive | Glutamic acid | 56-86-0 | C00025 | 84.04, 102.06, 130.05 |
| 156.08 | 16.29 | 1 | Positive | Histidine | 71-00-1 | C00135 | 110.07, 113.11, 156.08 |
| 127.07 | 11.18 | 1 | Positive | Imizadoleacetic acid | 645-65-8 |  | 81.04, 85.05 |
| 88.11 | 10.29 | 1 | Positive | Isoamylamine | 107-85-7 | C02640 | 88.08, 70.07 |
| 170.09 | 7.42 | 1 | Positive | Methyl-L-histidinate | 1499-46-3 |  | 110.07, 170.09 |
| 174.11 | 7.10 | 1 | Positive | N-Acetyl-L-leucine | 1188-21-2 | C02710 | 174.06 |
| 664.12 | 12.27 | 1 | Positive | NAD | 53-84-9 | C00003 | 428.04, 136.06, 524.06 |
| 124.04 | 8.08 | 1 | Positive | Niacin | 59-67-6 | C00253 | 124.04, 80.05, 96.04 |
| 166.09 | 10.11 | 1 | Positive | Phenylalanine | 63-91-2 | C00079 | 120.08, 166.09 |
| 166.07 | 9.82 | 1 | Positive | Phenylalanine | 63-91-2 | C00079 | 120.08 |
| 130.09 | 11.47 | 1 | Positive | Pipecolic Acid | 3105-95-1 | C00408 | 84.08, 130.09 |
| 218.14 | 9.86 | 1 | Positive | propionylcarnitine | 17298-37-2 | C03017 | 85.03, 218.14, 159.07 |
| 377.15 | 8.81 | 1 | Positive | Riboflavin | 83-88-5 | C00255 | 243.09 |
| 399.14 | 12.95 | 1 | Positive | S-Adenosyl-L-methionine | 29908-03-0 | C00019 | 250.09, 298.10, 136.06 |
| 106.05 | 13.17 | 1 | Positive | Serine | 56-45-1 | C00065 | 60.04 |
| 203.15 | 14.66 | 1 | Positive | Symmetric Dimethylarginine | 30344-00-4 |  | 203.15, 116.07, 70.07, 89.09 |
| 150.11 | 8.86 | 1 | Positive | Triethanolamine | 102-71-6 | C06771 | 70.07, 88.08, 132.10 |
| 183.09 | 12.33 | 1 | Positive | Triethyl phosphate | 78-40-0 |  | 98.98, 127.02 |
| 182.08 | 11.89 | 1 | Positive | Tyrosine | 60-18-4 | C00082 | 136.08, 123.04, 119.05, 91.05 |
| 296.08 | 7.13 | 1 | Negative | 5 -S- methyl -5- thioadenosine | 2457-80-9 | C00170 | 134.05 |
| 130.05 | 11.24 | 1 | Negative | 5-aminolevulinic acid | 106-60-5 | C00430 | 130.09 |
| 173.01 | 13.68 | 1 | Negative | Aconitic Acid | 585-84-2 | C00417 | 85.03, 111.01 |
| 505.99 | 12.85 | 1 | Negative | Adenosine triphosphate | 56-65-5 | C00002 | 505.99, 158.92, 408.01, 176.93 |
| 88.04 | 12.70 | 1 | Negative | Alanine | 302-72-7 | C01401 | 88.04 |
| 289.12 | 13.45 | 1 | Negative | Argininosuccinic acid | 2387-71-5 | C03406 | 131.08, 132.03, 173.1 |
| 132.03 | 12.85 | 1 | Negative | Aspartic acid | 56-84-8 | C00049 | 88.04, 71.01, 72.01 |
| 191.02 | 13.75 | 1 | Negative | Citric acid | 77-92-9 | C00158 | 87.01, 111.01, 85.03 |
| 221.06 | 13.56 | 1 | Negative | Cystathionine | 535-34-2 | C00542 | 134.03, 120.01, 221.06 |
| 147.03 | 12.76 | 1 | Negative | D-alpha-Hydroxyglutaric acid | 2889-31-8 | C01087 | 85.03, 101.02, 57.03, 103.04, 129.02 |
| 195.05 | 12.41 | 1 | Negative | D-Gluconic acid | 526-95-4 | C00257 | 75.01, 195.05, 129.02 |
| 171.10 | 7.09 | 1 | Negative | Decanoic acid | 334-48-5 | C01571 | 171.14 |
| 171.10 | 4.44 | 1 | Negative | Decanoic acid | 334-48-5 | C01571 | 171.14 |
| 171.14 | 4.16 | 1 | Negative | Decanoic acid | 334-48-5 | C01571 | 171.14 |
| 121.05 | 11.20 | 1 | Negative | Erythritol | 149-32-6 | C00503 | 71.01, 121.03, 89.02 |
| 259.02 | 13.43 | 1 | Negative | Glucose 6-phosphate | 56-73-5 | C00092 | 96.97, 78.96 |
| 129.02 | 12.76 | 1 | Negative | Glutaconic acid | 1724-02-3 | C02214 | 85.03, 101.02 |
| 146.05 | 12.70 | 1 | Negative | Glutamic acid | 56-86-0 | C00025 | 102.06, 128.03, 146.05 |
| 145.06 | 12.78 | 1 | Negative | Glutamine | 5959-95-5 | C00819 | 101.02, 145.06, 127.05 |
| 152.53 | 12.46 | 2 | Negative | Glutathione | 70-18-8 | C00051 | 74.02, 128.03, 152.87, 86.02 |
| 171.01 | 12.68 | 1 | Negative | Glycerol 3-phosphate | 57-03-4 | C03189 | 78.96, 96.97 |
| 171.08 | 11.65 | 1 | Negative | Glycylproline | 704-15-4 |  | 171.14, 96.96 |
| 521.98 | 13.97 | 1 | Negative | Guanosine triphosphate | 86-01-1 | C00044 | 521.98, 424.00 |
| 154.06 | 16.03 | 1 | Negative | Histidine | 150-35-6 | C00135 | 154.06, 137.03, 93.04 |
| 103.00 | 13.03 | 1 | Negative | Malonic acid | 141-82-2 | C00383 | 59.01, 73.03 |
| 181.07 | 13.89 | 1 | Negative | Mannitol | 69-65-8 | C00392 | 181.07, 101.02, 71.01 |
| 181.07 | 12.93 | 1 | Negative | Mannitol | 69-65-8 | C00392 | 181.07, 101.02, 71.01 |
| 181.07 | 14.25 | 1 | Negative | Mannitol | 69-65-8 | C00392 | 181.07, 101.02, 71.01 |
| 172.10 | 4.67 | 1 | Negative | N-Acetyl-L-leucine | 1188-21-2 | C02710 | 172.14, 130.09 |
| 190.05 | 4.93 | 1 | Negative | N-Acetyl-L-methionine | 65-82-7 | C02712 | 84.04 |
| 190.05 | 7.13 | 1 | Negative | N-Acetyl-L-methionine | 65-82-8 | C02713 | 84.04 |
| 122.02 | 8.09 | 1 | Negative | Nicotinic acid | 59-67-6 | C00253 | 94.03, 122.02, 78.03 |
| 164.07 | 10.12 | 1 | Negative | Phenylalanine | 63-91-2 | C00079 | 147.04, 164.07, 72.01 |
| 166.97 | 13.49 | 1 | Negative | Phosphoenolpyruvic acid | 138-38-9 | C00074 | 78.96 |
| 128.07 | 11.50 | 1 | Negative | Pipecolic acid | 535-75-1 | C00408 | 128.03 |
| 117.02 | 12.76 | 1 | Negative | Succinic acid | 110-15-6 | C00042 | 73.03, 117.02, 99.01 |
| 203.08 | 11.08 | 1 | Negative | Tryptophan | 54-12-6 | C00806 | 203.08, 116.05, 159.09 |
| 180.07 | 11.87 | 1 | Negative | Tyrosine | 60-18-4 | C00082 | 180.07, 163.04, 119.05 |
| 167.02 | 11.87 | 1 | Negative | Uric Acid | 13154-20-6 | C00366 | 69.01, 124.01, 96.02 |
| 151.03 | 11.20 | 1 | Negative | Xanthine | 69-89-6 | C00385 | 151.03, 108.02, 136.02 |

**Table S2.** Table of the 20 metabolites selected for targeted LC-MS analysis.

| m/z | Polarity | RT Start (min) | RT End (min) | Name | Abbreviation |
| --- | --- | --- | --- | --- | --- |
| 261.037 | Positive | 12.8 | 13.8 | Glucose 6-phosphate | G6P |
| 308.0911 | Positive | 11.8 | 12.8 | Glutathione reduced | GSH |
| 348.07036 | Positive | 11.5 | 12.5 | Adenosine monophosphate | AMP |
| 428.0367 | Positive | 11.9 | 12.9 | Adenosine diphosphate | ADP |
| 508.00304 | Positive | 12.4 | 13.4 | Adenosine triphosphate | ATP |
| 613.1593 | Positive | 12.8 | 13.8 | Glutathione oxidized | GSSG |
| 664.11641 | Positive | 11.1 | 12.1 | Nicotinamide adenine dinucleotide | NAD |
| 744.08272 | Positive | 12.4 | 13.4 | Nicotinamide adenine dinucleotide phosphate | NADP |
| 89.02441 | Negative | 9.5 | 10.5 | Lactic acid | Lac |
| 115.0037 | Negative | 12.5 | 13.5 | Fumaric acid | Fum |
| 117.01933 | Negative | 12.2 | 13.2 | Succinic acid | Succ |
| 133.0143 | Negative | 12.5 | 13.5 | Malic acid | Mal |
| 145.0143 | Negative | 12.1 | 13.1 | Alphaketoglutaric acid | a-KG |
| 166.9751 | Negative | 12.9 | 13.9 | Phosphoenolpyruvic acid | PEP |
| 168.9908 | Negative | 12.2 | 13.2 | DHAP | DHAP |
| 191.0197 | Negative | 13.1 | 14.1 | (Iso)citrate | (Iso)cit |
| 229.0119 | Negative | 12.4 | 13.4 | Ribose 5-phosphate/Ribulose 5-phosphate | R5P/Ru5P |
| 259.0224 | Negative | 12.8 | 13.8 | Fructose 6-phosphate | F6P |
| 289.033 | Negative | 12.5 | 13.5 | Sedoheptulose 7-phosphsate | SH7P |
| 338.9888 | Negative | 13 | 14 | Fructose 1,6-biphosphate | FBP |

